# A Research-Integrated Curriculum on CAR-T Cell Therapy

**DOI:** 10.64898/2026.09.23.753877

**Authors:** Youan H. Khan, Lauren Rice, Melissa C. Srougi

**Affiliations:** Biotechnology Program and, North Carolina State University Raleigh, NC United States; Department of Molecular Biomedical Sciences, North Carolina State University Raleigh, NC United States

## Abstract

Chimeric antigen receptor (CAR) T cells are patient-derived T cells engineered to recognize specific antigens on malignant cells, representing a major breakthrough in personalized cancer immunotherapy. While CAR-T cell therapy is an established treatment for relapsed or refractory hematologic malignancies, understanding the rationale and methodology behind its development is essential for training the next generation of oncology researchers. Here, we describe a novel, hands-on small-enrollment laboratory course designed to teach upper-level undergraduate and graduate students about immunotherapies and CAR-T cell technology by paralleling a typical bench-scale development process. Students isolated T cells from porcine peripheral blood mononuclear cells (PBMCs) via immunomagnetic selection, followed by flow cytometry to assess purity, viability, and yield. Students then used pre-engineered B7H3 CAR-T cells to perform co-culture assays with U87 glioblastoma cells to evaluate tumor cell killing efficacy. Cytotoxicity was qualitatively evaluated using immunofluorescence microscopy, while IFN-γ release was quantified using Enzyme-linked Immunosorbent Assay (ELISA). Student performance was evaluated through electronic laboratory notebooks, discussion questions, a capstone project, and student-generated data. Our results demonstrate students’ high academic performance across assignments and the successful execution of complex laboratory workflows. Collectively, these findings establish the feasibility and implementation of a research-integrated curriculum providing foundational exposure to immuno-oncology research tools and CAR-T cell production techniques.

## Introduction

Recent methods in cancer immunotherapy leverage the human immune system to develop highly selective therapeutic agents against a spectrum of malignancies [1]. Among these approaches, autologous T cells engineered with chimeric antigen receptors (CAR) have emerged as a powerful treatment modality. CAR-T cells enhance targeted cancer cytotoxicity and persistent immune response, while bypassing MHC-restricted antigen presentation. Consequently, CAR-T cell therapies have demonstrated significant clinical efficacy against several refractory or relapsed hematological malignancies [2-5].

To date, seven therapies have received FDA-approval for indications including B-cell acute lymphoblastic leukemia, follicular lymphoma, large B-cell lymphoma, multiple myeloma, and mantle cell lymphoma [6, 7]. While CAR-T therapies have achieved success in treating hematologic malignancies, including long-term survival in some patients [4, 5], significant barriers still exist in their application to treating solid tumors [7]. Overcoming these barriers requires research focused on discovering novel receptor targets, enhancing construct potency [2], and developing combination therapies [3, 7].

As CAR-T cell therapies continue to be developed and applied in clinical practice, it is important to ensure that future cancer biologists have the foundational knowledge and skills necessary to bring these revolutionary treatments from research, to market, and to clinical practice. Initiatives in furthering cancer education have successfully introduced undergraduate and graduate students to cancer genetics and bioinformatics [8-10], and several seminar-based, interdisciplinary immuno-oncology courses exist at the graduate level [11, 12]. However, few offer practical hands-on laboratory experiences, particularly in the cell therapy sphere. Early research exposure is critical for students and has been shown to significantly enhance student persistence in STEM and cancer-related-research [13-16]. Since not all students are able to participate in independent research, course-based undergraduate research experiences (CUREs) can address this gap by providing early experiential training [17, 18], fostering scientific confidence [19, 20], strengthening their scientific identity [21] and preparing undergraduate students for careers in cell and gene therapeutics. Bridging theoretical tumor immunology with practical cellular cancer-related laboratory experiences requires curricula that explicitly integrate core cancer biology concepts with laboratory application.

To fulfill this need, we developed an 8-week bench-scale laboratory curriculum centered on CAR-T cell therapy. The course covers key techniques required to produce CAR-T cells, including T cell isolation, flow cytometric quality control, CAR expression assessment, and target cell-killing assays. The curriculum was intentionally designed using the CURE framework, which emphasizes collaboration, discovery, relevance, and iteration [20]. Specifically, collaboration was fostered through paired lab partners in real-world workflows for the hands-on execution of lab techniques and team capstone projects; relevance was established by targeting B7H3 solid tumor antigens; discovery was established through student-driven experiments evaluating cell killing efficacy; and iteration was embedded into the flow cytometry and ELISA labs, data analysis, and structured reflection. Here, we describe the design, implementation, and feasibility of this research-integrated course and evaluate overall student academic performance.

## Methods

### Participants and Course Context

The laboratory was successfully implemented by students enrolled in a dual 400/500 level 8-week lecture and laboratory course called Cell Therapies at NC State University. This is an interdisciplinary STEM course that discusses biotechnology, biomanufacturing processes of cell-based cancer immunotherapies. While the lecture reviews the principles and types of cell therapies, the laboratory focuses on the bench-scale biotechnology of CAR-T cell therapy production and immunotherapies in oncology. Students enrolled in this course were 4th-year undergraduate students and 1st year graduate students majoring in a variety of areas including but not limited to comparative biomedical sciences, chemical engineering, and biomanufacturing **(Supplementary Fig 1)**. A total of 12 students were enrolled in the course, which is offered once per academic year. The course includes a weekly 2.5-hour lecture period and a 5-hour laboratory. Prerequisites for enrollment in the course include two semesters of organic chemistry, and a foundational course on the manipulation of recombinant DNA or equivalent course.

### Laboratory Curriculum

Laboratory experiments followed key steps in the biomanufacturing of a CAR-T cell therapy at bench scale (**Fig 1**). All experiments were performed *in vitro* using mammalian cell culture. Students began by isolating primary T cells from porcine peripheral blood mononuclear cells (PBMCs) by immunomagnetic positive selection. The efficacy and viability of the T cell isolation was validated via flow cytometry. Next, students were provided with activated human T cells transduced with B7H3-CAR and evaluated cell transduction efficiency and viability via flow cytometry. Following validation of CAR expression, students performed co-culture cell killing assays with B7H3 and green fluorescent protein (GFP) expressing human U87 glioblastoma cells. Students plated cells at various effector to target ratios and evaluated the effectiveness of cell-killing by measuring human IFN-γ release via ELISA assay and qualitatively by observing GFP fluorescence of U87 cells with immunofluorescence microscopy. Students were given detailed protocols and were required to complete pre-lab entries prior to lab in which they summarized the laboratory’s purpose and wrote out detailed methods. The course syllabus outlined the student learning outcomes and conceptual and technical skills as well as troubleshooting techniques.

**Fig 1.**
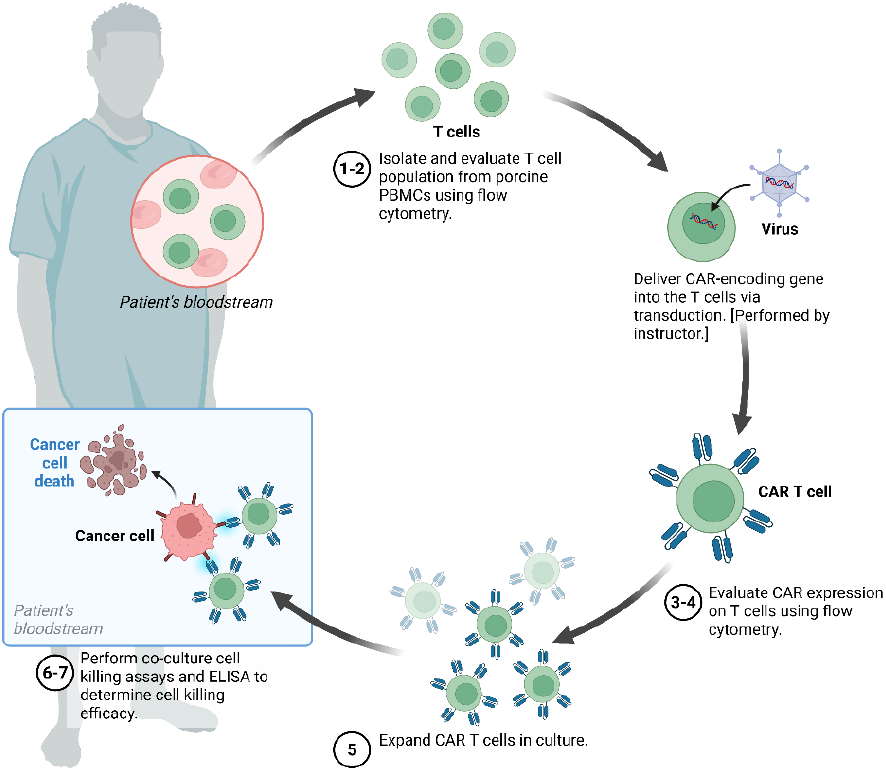
Diagram of classroom laboratory experiments for the bench scale manufacturing of B7H3 CAR-T cells. Prior to the course, students practice sterile mammalian cell culture techniques. (Labs 1-2) Students isolate primary T cells from porcine PBMCs via immunomagnetic positive selection and evaluate isolation efficacy and viability via flow cytometry. Instructor transduces primary human T cells with B7H3 CAR construct. (Labs 3-4) Students analyze B7H3 CAR surface expression and cell viability using flow cytometry. (Lab 5) Students expand CAR-T cells in culture. (Labs 6-7) Finally, students perform co-culture cell-killing assays and quantify human IFN-γ secretion via ELISA to evaluate CAR-T cell cytotoxicity against target glioblastoma cells. Image Created in BioRender. (2026) https://BioRender.com/pzo2d5y

The specific student learning outcomes (SLOs) for the laboratories are listed below. By the end of the course, students will be able to:

1. Understand the basic physiology of immune cell types, MHC classes, and immune responses.
2. Describe the biological principles and mechanisms of action of major cell therapies, including CAR-T cell and stem cell-based treatments.
3. Summarize the common biomanufacturing processes that create CAR-T cell-based products.
4. Apply core aseptic techniques and standard operating procedures to successfully isolate, culture, genetically engineer, expand, and maintain mammalian cell lines for cell therapy use.
5. Interpret and analyze data from laboratory experiments to draw valid scientific conclusions about CAR-T cell biology and efficacy.
6. Evaluate the efficacy, safety, and current challenges of cell therapy products in oncological practice by interpreting data from scientific literature.
7. Reflect on their own thinking and the thinking of others from a variety of disciplines and backgrounds.

### Course Assessment Methods

To evaluate student academic performance without relying solely on final letter grades, student work was evaluated through multiple assessments mapped directly to specific student learning outcomes (SLOs; **Supplementary Table 1**). Structured rubrics were utilized to measure technical lab execution, data analysis accuracy, reflective depth, and synthesis of information (**Supplementary Tables 1-3**). While high assessment scores reflect thorough coursework execution rather than pre/post learning gains, this alignment ensured that student evaluations directly corresponded to conceptual and technical course goals. The primary assessment was students’ electronic laboratory notebooks (ELNs; LabArchives software), which included detailed purpose, methods, results, and discussion. The lab notebooks were used to assess student learning outcomes (SLOs) (1-7). Students’ responses to discussion questions after each lab were used to determine their critical analyses and understanding of their individual data (SLOs 4-7). Furthermore, students were required to complete reflections related to the techniques and concepts learned as part of the course, which was used to assess their ability for critical reflection (SLO 7). Although students worked in pairs in the laboratory, the ELN entries, discussion questions and reflection prompts were individual efforts. Throughout the half-semester, students worked collaboratively in teams of 3-4 on a cumulative capstone project. Students selected emerging topics in cellular therapies to research and present (SLOs 1-3, 5-7). The capstone project required students to synthesize both conceptual and technical knowledge, evaluate peer-reviewed literature, reflect on ethical dilemmas associated with modern translational therapies, and present their findings to a scientific audience using a format of their choosing. Details regarding assessment rubrics, grading criteria and associated SLOs are found in **Supplementary Tables 1-3**.

This study was reviewed and determined to be exempt by the NC State University Institutional Review Board (IRB #28928).

## Results

Participants (n=12) were students enrolled in a dual upper-level cell therapies course at an R1 University in the southeastern United States. Participants were comprised of 11 undergraduate students (91.7%) and 1 graduate student (8.3%) across multiple academic majors, including: Chemical Engineering (n= 5; 41.7%), Biochemistry (n=3; 25%), and 1 student from each of the following major programs: Microbiology (8.3%), Textile Technology (8.3%), Genetics (8.3%), and Comparative Biomedical Sciences (8.3%) (**Supplementary Figure 1**).

Student academic performance was evaluated using electronic lab notebooks (ELNs), in-lab discussion questions, written reflections, and student generated data. These assessments were evaluated based on student technical execution of experiments, data analysis, content accuracy, reflective depth, and other metrics which are detailed in the provided rubrics **(Supplementary Tables 1-3)**. Weekly ELN submissions averaged 99.3% ± 1.3% across all completed assignments (**Fig 2**). On post-lab discussion questions evaluating data synthesis and biological concepts (SLOs 4-7), students averaged 98.0% ± 1.5% (**Fig 2**). On the written reflection assignments which evaluated student comprehension of bench-scale laboratory workflows and their alignment with current industry therapeutic manufacturing standards (SLO 7), students (n=11) averaged 93.9% ± 8.0% (**Fig 2**). Finally, on the cumulative team capstone project, student teams (n= 12) achieved an average score of 90.8% ± 3.9% (**Fig 2**).

**Fig 2.**
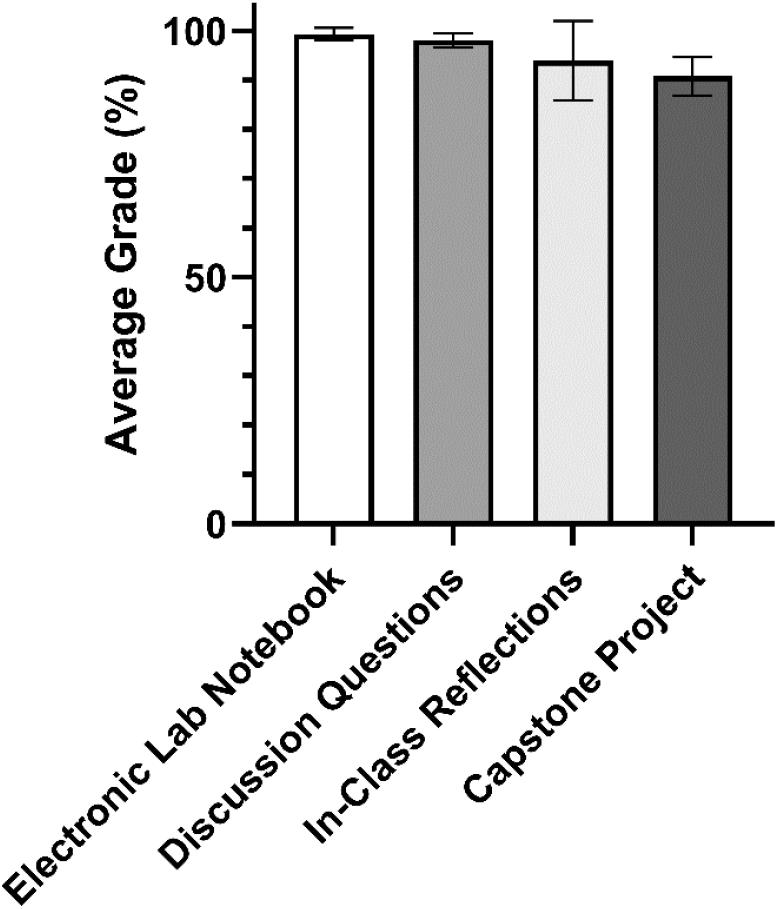
Average student performance across course assessment types. Mean student grades on laboratory-related assignments, including Electronic Lab Notebook (ELN) submissions (n = 12), discussion questions (n = 12), reflection prompts (n = 11), and the collaborative capstone project (n = 12). Data are presented as mean +/-SD. Unsubmitted assignments were excluded from final statistical analyses (one reflection prompt assignment was unsubmitted and omitted, n = 11).

Representative examples of student work demonstrating the successful execution of each stage in the bench-scale CAR-T cell biomanufacturing workflow (**Fig 1**) are shown in **Figure 3A-H**. Most students successfully isolated primary T cells from porcine PBMCs and prepared these samples for downstream flow cytometric analysis (**Fig 3A**). Next, students applied flow cytometry to validate B7H3 CAR expression within the primary human T cell populations and confirmed CAR expression (**Fig 3B**). The full gating strategy is found in **Supplementary Figures 2 and 3**. To evaluate CAR-T cell functional cytotoxicity, students performed immunofluorescence (IF) microscopy in co-culture with GFP-expressing human U87 glioblastoma cells across various effector-to-target (E:T) ratios (**Fig 3C-G**). Student data demonstrated a dose-dependent reduction in GFP fluorescence at higher E:T ratios (10:1, 5:1, and 1:1), indicating effective CAR-T cell-mediated tumor killing (**Fig 3C-F**) that was absent in the activated T cell control (**Fig 3G**). This qualitative assessment was supported by quantitative analysis of CAR-T cell activation, measured by human IFN-γ secretion using ELISA (**Fig 3H**). A time-dependent increase in IFN-γ release was observed at E:T ratios 10:1, 5:1, and 1:1 (**Fig 3C-F**), whereas minimal cytokine release was detected at the 1:10 E:T ratio and in control (**Fig 3H**). Collectively, these representative laboratory results demonstrate successful student execution of complex CAR-T laboratory protocols (**Fig 3**).

**Fig 3.**
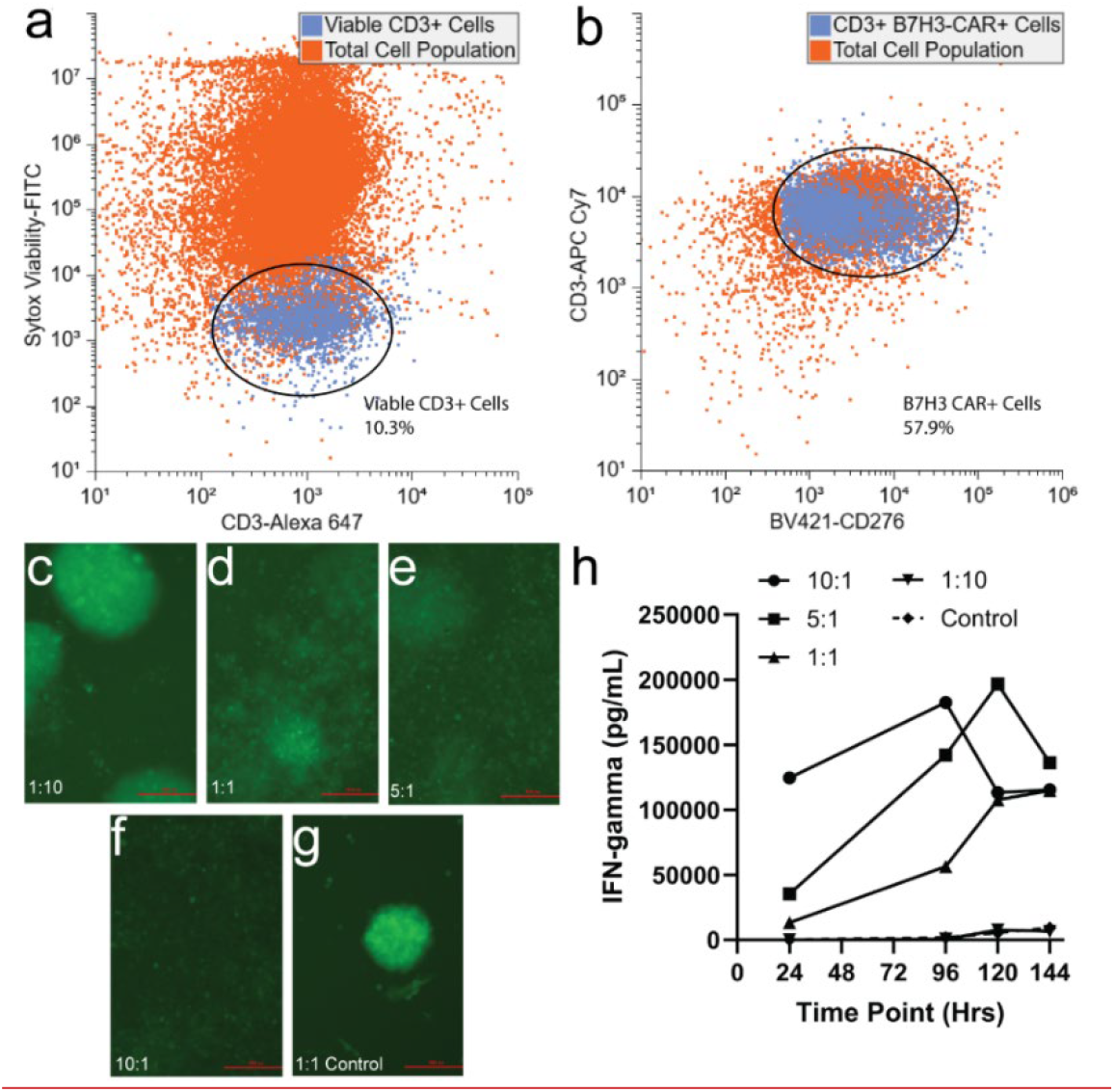
Representative student-generated experimental data across the CAR-T biomanufacturing workflow. **(A)** Flow cytometric purity and viability analysis of CD3+ primary T cells isolated via immunomagnetic selection from porcine peripheral blood mononuclear cells (PBMCs). Viability was determined using SYTOX dead cell stain. **(B)** Representative gating strategy evaluating B7H3 CAR expression post T-cell transduction. Flow cytometry data were standardized and re-gated to maintain anonymity and consistency across cohorts. **(C–G)** Immunofluorescence (IF) microscopy of GFP-expressing U87 glioblastoma cells after co-culture with B7H3 CAR-T cells at 120 hours across effector-to-target (E:T) ratios: **(C)** 1:10, **(D)** 1:1, **(E)** 5:1, **(F)** 10:1, and **(G)** an activated, unmodified T cell control (1:1 E:T). Images taken with a Nikon Fluorescent Eclipse Ts2 with 10x objective. Scale Bar 200 µm. **(H)** Quantification of IFN-γ secretion in co-culture supernatants measured via ELISA and performed in duplicate over time 24-144 h.

## Discussion

This study evaluates a novel laboratory curriculum focused on the generation of CAR-T cells. CAR-T therapies have emerged as a lifesaving intervention for a variety of malignancies, with an expanding therapeutic scope driven by ongoing efforts to enhance their efficacy, safety, and accessibility [3, 4, 6, 7]. Hands-on training in the foundational techniques and principles of cell therapy provides students with practical exposure to core concepts in cancer therapy and immuno-oncology that are difficult to convey in existing lecture-only courses. Moreover, to meet the growing demand for skilled personnel in cellular therapeutics manufacturing, it is imperative that future biotechnologists receive practical hands-on training in the foundational techniques and principles of cell therapy production [22].

The curriculum described here enables an interdisciplinary student cohort to explore CAR-T cell design and bioprocessing across all production stages (**Fig 1)**, leveraging accessible, bench-scale biotechnology platforms while learning about core immunotherapy concepts. The course enrolled students from diverse academic majors (**Supplementary Fig 1**), reflecting the inherently interdisciplinary nature of cancer biology and cell therapy research. While the labs were conducted with a small cohort of students, the laboratory protocols are scalable for larger courses and utilize commercially available materials, such as porcine-derived blood cells, immunomagnetic selection kits, and standard ELISAs.

To connect coursework with cutting-edge developments in solid tumor immunotherapy, laboratory exercises centered on evaluating a CAR construct targeting the B7H3 antigen [23, 24]. By using clinically relevant B7H3-targeted CAR against U87 glioblastoma cells, students directly engaged with concepts central to cancer education in solid tumor immunotherapy, including tumor antigen selection, the immunosuppressive glioblastoma tumor microenvironment, and the off-target limitations that govern CAR-T efficacy against solid tumors [7, 24]. The laboratory course described allowed students to connect fundamental cancer biology concepts, with the principles of precision oncology that guide antigen and construct selection in cell therapy development [23]. Importantly, these protocols can be tailored to alternative established or emerging tumor targets. The core competencies learned in this course, including aseptic technique, sterile mammalian cell culture, and quality control assays, remain directly transferable across broad oncology sectors.

The CURE pedagogical framework fosters a high-impact experiential learning experience [19] in which undergraduate and graduate students take ownership of their laboratory experiments. This modular approach ensures that the course remains adaptable to evolving cell therapy modalities and manufacturing pipelines. Student performance data demonstrate that participants earned high average grades across all assessment types (**Fig 2**). In addition to theoretical concepts, students in this small cohort successfully executed complex laboratory workflows, generating data demonstrating effective T cell isolation, CAR expression, and functional *in vitro* cell killing efficacy (**Fig 3 and Supplementary Figures 2-3**).

## Limitations

Several limitations of this study should be noted. First, the evaluation was conducted with a small cohort (n=12) of upper-level undergraduate and graduate students at a single research-intensive university. Because course enrollment was elective rather than a degree requirement, self-selection bias and class size may limit the direct generalizability of these findings to a broader student population. High assignment performance in this self-selected cohort may partially reflect baseline academic motivation and prerequisite preparation rather than curriculum-driven gains alone [25]. Second, this study did not include a pre/post comparative assessment instrument of student knowledge. Therefore, the reported assessment scores may reflect successful completion of coursework and alignment with rubric criteria rather than direct learning gains.

## Conclusions

This study demonstrates the feasibility of a research-integrated, bench-scale CAR-T cell laboratory curriculum designed for an interdisciplinary university student cohort. By bridging foundational oncology concepts with hands-on laboratory techniques, the course modeled translational cancer research workflows within an academic setting. Students demonstrated strong academic performance across diverse assessments while successfully executing critical procedures in CAR-T cell engineering, quality control, and cytotoxicity evaluation. Although implemented at bench scale without automated systems or clinical-grade infrastructure, the modular nature of this research-integrated framework allows for straightforward adaptation across diverse institutional settings and emerging target tumor antigens. Ultimately, embedding scalable cancer biology laboratory experiences into higher education offers a promising, adaptable model for providing undergraduate and graduate students with hands-on exposure to the technical competencies underlying immunotherapies in cellular oncology.

## Acknowledgements

The authors would like to thank the students enrolled in the course for their feedback and participation. We gratefully acknowledge the NC State Provost’s Office, the Biotechnology Program, and the Biomanufacturing Teaching and Education Center for their generous support, resources, and facilities. We also thank Dr. Yevgeny Brudno and Dr. Madelyn VanBlunk for their guidance during course development and for generously providing reagents as well as Dr. Elisa Crisi for providing porcine PBMCs.

## Declarations Financial

No funding was received for conducting this study. The authors have no relevant financial or non-financial interests to disclose.

