## Supplementary Information for "A Research-Integrated Curriculum on CAR-T Cell Therapy"

| **Student Learning Outcome (SLO)** | **Assessment** | **Measurements** | **Rubric Scoring Criteria** |
|---|---|---|---|
| SLO 1-3 Theoretical concepts of immune physiology, CAR T mechanisms, and lab techniques. | • Capstone project<br>• Electronic laboratory notebooks<br>• Discussion questions | Synthesis of literature on emerging therapies, presenting experimental pipelines, and explaining target biological mechanism(s) of action. | Depth of biological rationale, accuracy, and integration of primary peer-reviewed literature. |
| SLO 4 Application of aseptic technique, T-cell isolation, and cell culture maintenance. | • Laboratory generated data<br>• Electronic laboratory notebooks<br>• Instructor observation | Documentation of the step-by-step standard operating procedure, reporting experimental data including cell yields, purity, viability and capturing/ gating flow cytometry data. | Complete protocol execution, accurate data recording, accurate gating strategies, viability analyses, ability to isolate T cells from PBMCs, ability to culture primary cells long-term (multiple weeks) and maintain contamination free. |
| SLO 5 & 6 Data interpretation and literature evaluation. | • Electronic laboratory notebooks<br>• Discussion questions<br>• Capstone project | Analyzing T cell isolation, viability, and purity; analyzing CAR T transduction efficiency and viability; co-culture immunofluorescence data of co-culture cell killing assays, ELISA IFN-gamma concentrations against a standard curve. Discussing results in the greater context of the field. | Accurate experimental calculations, statistical analyses and interpretation of experimental outcomes, evaluation of limitations and suggestions for improvement. |
| SLO 7 Reflective thinking across interdisciplinary backgrounds. | • Electronic laboratory notebooks<br>• Discussion questions<br>• Capstone project | Responding to prompts and identification of ethical considerations, translational bottlenecks, cross-disciplinary collaboration. | Critical self-reflection, balanced articulation of the pros vs cons of interdisciplinary perspectives, connection to current industry standards. |

**Supplementary Table 1.** Student learning outcomes alignment with course assessment methods and measures.

| Criteria | Exemplary (100%) | Proficient (80-89%) | Developing (60-79%) | Missing (0-59%) | Weight % |
|---|---|---|---|---|---|
| Title | Unique descriptive title in own words, data and initials. Correct Lab Archives page setup. | Descriptive title, but missing minor details (data, initials omitted) | Title is a repeat of generic protocol; missing multiple data fields | Title missing or entry incorrectly structured. | 5 |
| Purpose | Concise 2-3 sentence purpose in own words. Explicitly names methods and connects them directly to broader laboratory goals. | Correct length and details methods, but weakly articulates connection to broader field and lab goals. | Exceeds length, restates protocol steps without naming core methods or overall course goals. | Missing, copied verbatim from protocol, or fails to define scientific goals. | 5 |
| Methods & Experimental Execution | Meticulously records all methods and real-time modifications, incubation variations, volumes, or aseptic deviations. | Logs major procedural modifications, but misses minor execution details or deviations. | Fails to document actual changes/deviations made during experimentation. | Protocol is missing or incomplete; no real-time documentation provided. | 20 |
| Results (Data & Annotation) | High-quality raw data provided. Fully annotated (e.g. gel lane labels, standards, flow axes, cell counts, viability %). | Raw data present with basic annotations; minor labeling gaps. | Data present but poorly annotated, blurry, inaccurate, or unreadable. Unable to discern experimental conditions. | Raw data missing or incomplete, unannotated, or improperly borrowed without authorization. | 15 |
| Results (Data Narrative) | Comprehensive, self-contained text referencing all figures/tables. Accurately describes data trends so they are visible and understandable. | Narrative references figures/tables correctly, but lacks sufficient description to stand independently. | Vague or overly brief narrative, fails to references specific visuals or merely lists raw values. | Text narrative is missing, or directly contradicts the visual data presented. | 15 |
| Discussion and Critical Reflection | Data is analyzed thoroughly and with scientific accuracy. Draws connections with the scientific literature and conveys interdisciplinary thinking. Troubleshoots experiments that do not work. | Lacks critical depth in analysis, literature context, or complete analysis. Provides superficial explanations for experiments that do not work. | Discussion is partially addressed, superficial, or skips multi-part requirements. Reflection is not thorough. Provides inaccurate or incomplete explanations for experiments that do not work. | Discussion section omitted or majority of questions unanswered. No reflection provided. No troubleshooting or improvement provided. | 40 |

**Supplementary Table 2.** Electronic notebook rubric, which includes integrated discussion and reflection questions.

| Category | Absent | Inadequate | Developing | Good/Excellent | Score (out of 60 max points) |
|---|---|---|---|---|---|
| Scientific content and accuracy | 0 points No explanation. | 1-5 points Some explanation but mostly superficial. Explanation was not scientifically accurate. Level of explanation of content is not appropriate for the scientific audience. | 6-7 points Partially explained but missed some important details. Level of explanation is partially suitable for target audience. | 8-10 points Thorough explanation of the scientific background on the topic. Mechanism(s) are detailed and accurate. | 10 |
| Professionalism and ability to answer questions. | 0 points Needs work. | 1-5 points Incomplete explanation to questions and/or informal style. | 6-7 points Serious style but unable to answer questions accurately based on presented materials. | 8-10 points Professional and clearly articulates answers to questions. Demonstrates the ability to connect the topic to the broader immunology or biotechnology/engineering principles. | 10 |
| Trend analysis | 0 points Did not discuss or discussion is inaccurate. | 1-5 points Topic is dated (e.g. basic 1st gen CARs) with little emerging context. | 6-7 points Identifies trends in the last 3 years and explains its basic clinical relevance. | 8-10 points Critically evaluates the trend against current "Gold Standards." Identifies and discusses specific gaps it fills. | 10 |

| Category | Absent | Inadequate | Developing | Good/Excellent | Score (out of 60 max points) |
|---|---|---|---|---|---|
| Communication (organization, visuals, data elocution and creativity) | 0 points Disorganized and difficult to follow. Data is copied without clear interpretation | 1-5 points Needs work. Artifact visually cluttered (text/images) and/or not clearly communicated verbally/written such that it was hard to understand what was going on. | 6-7 points Clean visuals; data supports the main narrative. Organized but not explained fully or well. | 8-10 points High level visuals. Explain concepts clearly and thoroughly. Text, visuals, verbal communication was clear and organized. Easy to follow. | 10 |
| Translation to clinic | 0 points Does not address how the therapy moves from the lab to the patient. | 1-5 points Some mention of translational medicine perspectives but are vague, general and/or inaccurate for the technology. | 6-7 points Discusses general hurdles like cost or FDA approval or clinical trial. | 8-10 points Provides a detailed “bench to bedside” plan, addressing specific chemistry, manufacturing and other controls. How it will overcome common clinical hurdles such as tumor microenvironment or other. | 10 |

| Category | Absent | Inadequate | Developing | Good/Excellent | Score (out of 60 max points) |
|---|---|---|---|---|---|
| Bioethics | 0 points Does not discuss bioethical dilemmas or discusses issues that are not relevant. | 1-5 points Ideation employed creative thoughts but artifact(s) was not creative. Likely would not draw audience attention. | 6-7 points Overall presentation shows some level of creativity, and audience interest leading to passable topic/artifact that falls somewhat short on detail. | 8-10 points Overall shows an acceptable level of creativity that would draw audience attention leading to a satisfactory product that can be used for educational purposes. | 10 |
| References | Up to -10 pts may be deducted if students do not have a minimum of 8 references that are appropriately cited. Students are responsible for vetting these references. | | | Total Score | 60 |

**Supplementary Table 3.** Capstone project rubric.

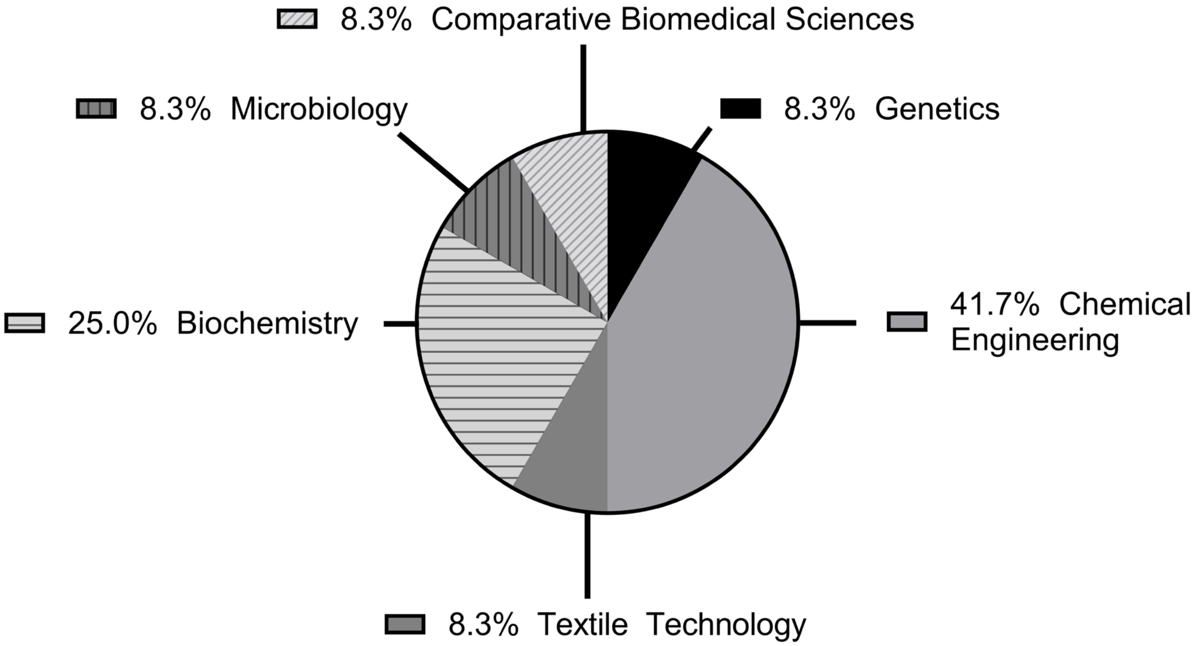


**Supplementary Figure 1.** Class majors for students (n=12) enrolled in the Cell Therapies course.

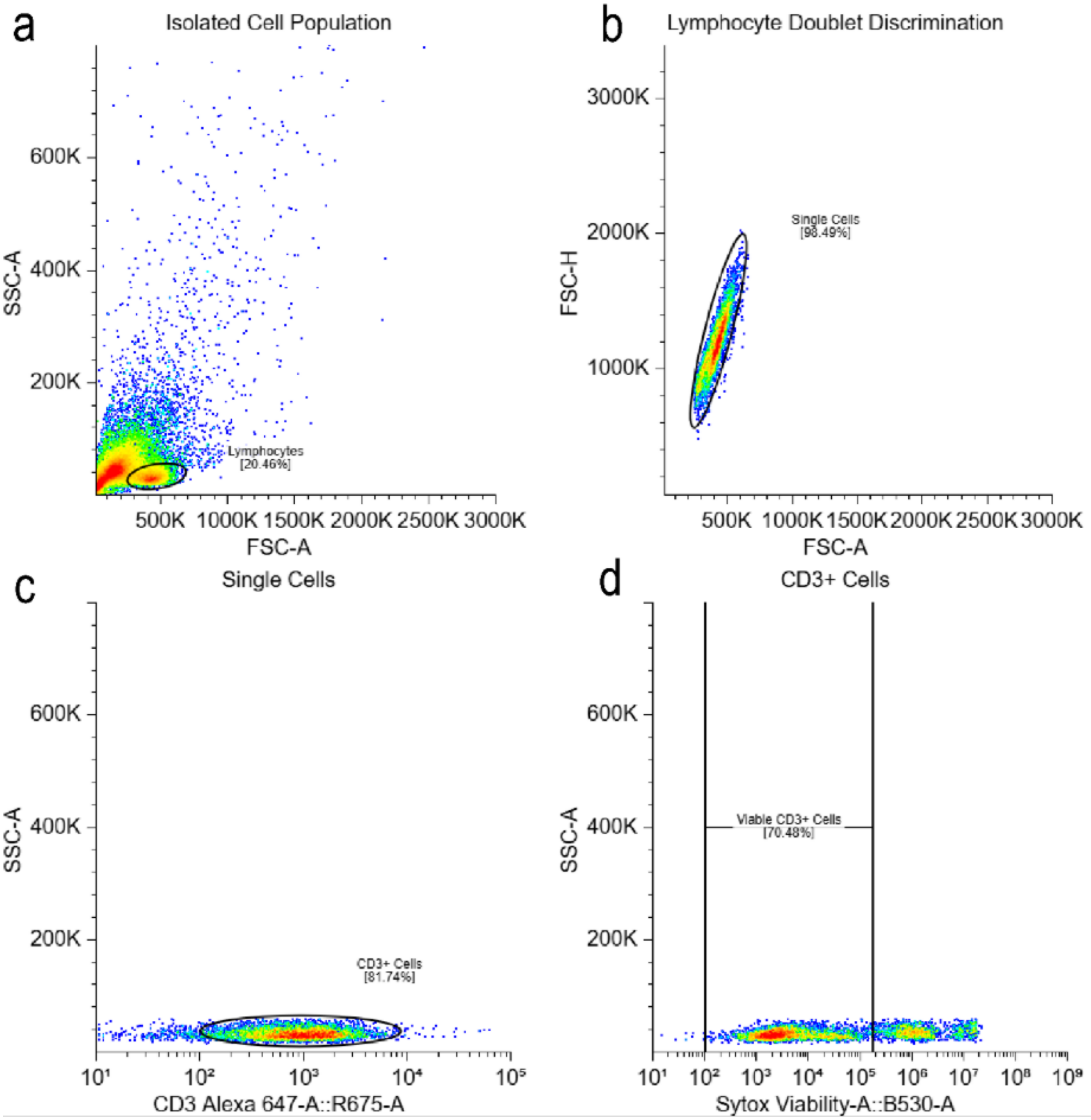


**Supplementary Figure 2. Student flow cytometric analysis of immunomagnetically isolated T cells.** Representative gating strategy used by students (analyzed via Floreada software) to evaluate T-cell purity and viability following positive selection from porcine peripheral blood mononuclear cells (PBMCs). Sequential gating displays the **(a)** total lymphocyte population, **(b)** doublet discrimination, **(c)** CD3+ surface marker expression, and **(d)** viable CD3+ T cells. Subpopulation percentages are indicated within each gate.

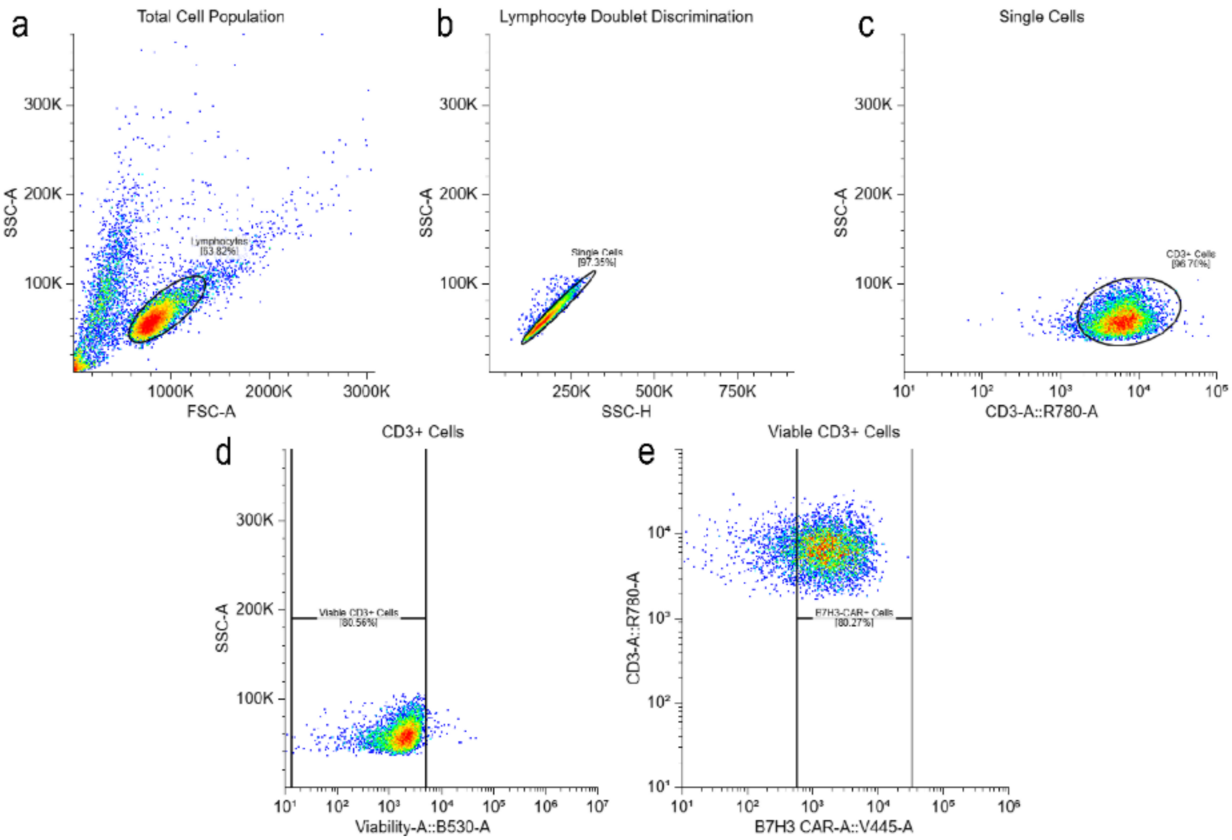


**Supplementary Figure 3. Student flow cytometric analysis of transduced B7H3-CAR T cells.** Representative gating strategy executed by students using Floreada software to evaluate transduction efficiency and viability in primary human T cells. Sequential gating displays the **(a)** total lymphocyte population, **(b)** doublet discrimination, **(c)** CD3+ T-cell selection, **(d)** viable CD3+ cells, **(e)** and surface expression of the B7H3 CAR. Population percentages are indicated within each gate.
